# Single-cell transcriptome profiling reveals dermal and epithelium cell fate decisions during embryonic hair follicle development

**DOI:** 10.1101/704379

**Authors:** Wei Ge, Shao-Jing Tan, Shan-He Wang, Lan Li, Xiao-Feng Sun, Wei Shen, Xin Wang

## Abstract

Characterization of the morphological structure during hair follicle development has been well documented, while the current understanding of the molecular mechanisms involved in follicle development remain limited. Here, using unbiased single-cell RNA sequencing, we analyzed 15,086 single cell transcriptome profiles from E13.5 and E16.5 fetal mice, and newborn mouse (postnatal day 0, P0) dorsal skin cells. Based on t-distributed Stochastic Neighbor Embedding (tSNE) clustering, we identified 14 cell clusters from skin cells and delineated their cell identity gene expression profiles. Pseudotime ordering analysis successfully constructed epithelium/dermal cell lineage differentiation trajectory and revealed sequential activation of key regulons involved during cell fate decisions. Along with this, intercellular communication between different cell populations were inferred based on a priori knowledge of ligand-receptor pairs. Together, our findings here provide a molecular landscape during hair follicle epithelium/dermal cell lineage fate decisions, and more importantly, recapitulate sequential activation of core regulatory transcriptional factors for different cell populations during hair follicle morphogenesis.

## Introduction

During hair follicle development the dynamic morphological changes have been extensively studied (Paus et al., 1999; Saxena et al., 2019a; Schmidt-Ullrich and Paus, 2005). In mice, *in utero* hair follicle development has been histologically categorized into three unique stages: induction (E13.5 - E14.5), organogenesis (E15.5 - 17.5), and cytodifferentiation (E18.5 onwards) (Schmidt-Ullrich and Paus, 2005). More recently, with the development of single-cell RNA sequencing (scRNA seq), new intermediate cell states during early hair follicle morphogenesis have been delineated and an updated classification of different hair follicle stages has been reported (Mok et al., 2019; Saxena et al., 2019a). Seminal works have delineated that reciprocal signaling pathways between epithelial and dermal cell populations play vital roles during hair follicle morphogenesis (Lee and Tumbar, 2012; Millar, 2002; Rishikaysh et al., 2014; Sennett and Rendl, 2012). However, our current knowledge regarding in utero hair follicle morphogenesis remains limited.

At about E13.5 in mice, the unspecified epidermis received signals from mesenchyme (also known as “first dermal signal”) and subsequently forms a layer of thickening epithelial known as placodes, seen as the earliest morphological characteristic marking the initiation of hair follicle morphogenesis (Biggs and Mikkola, 2014; Hardy, 1992). Wnt/β-Catenin and Eda/Edar/NF-κB signaling have been demonstrated to play vital roles during this earliest placode fate commitment (Chen et al., 2012; Schmidt-Ullrich et al., 2006), while the upstream regulators remain elusive. Following placode fate commitment, placodes signal to underlying fibroblast to promote the formation of dermal condensate (DC), the precursor of the dermal papilla (DP). The signal/s involved in this “first epithelial signal” remain little known. However, fibroblast growth factor 20 (Fgf20) signaling has been shown to be one of the “first epithelial signals” as ablation of Fgf20 in mice results in the failure of DC formation (Biggs et al., 2018). After the commitment of the placode and DC, the cross talk between placode and DC then promotes transition to the next stage of development: signals from DC, also known as the “second dermal signal”, promote the specification of the upper placode into hair follicle stem cell (HFSC) precursors or lower layer matrix precursors (Ouspenskaia et al., 2016), during which Wnt and Shh signaling has been demonstrated to play a role (Ouspenskaia et al., 2016). As development continues, the DC signal to the upper epithelium and promote the downward proliferation of the epithelium into the dermal layer, and finally, result in the envelopment of the DC, with Shh and Pdgfa signaling demonstrated to participate in this process (Chiang et al., 1999; Karlsson et al., 1999). After the envelopment of the DC by epithelial cells, the DC at this stage then matures into the DP surrounded with matrix cell populations. As the cross talk between the DP and surrounding matrix continues, signals from the DP then promote the surrounding matrix cells to further differentiate into the hair shaft and inner root sheath (IRS), during which the molecular signature has been sparsely reported. At this time, the rudiment of a developed hair follicle becomes morphologically obvious.

While the process of hair follicle morphogenesis has been well-documented, during hair follicle development, our current understanding of the molecular signatures and gene regulatory networks operating within a particular cell population remains limited. Besides, limited progress has been made to identify conserved gene markers expressed present in different cell populations. By using genetic loss-of-function assays, transgenic mouse models, and flow cytometer cell sorting technology based on prior knowledge of well-defined markers, the molecular signatures of different cell populations during hair follicle development has been reported (Driskell et al., 2009; Mok et al., 2019; Rezza et al., 2016). However, the molecular signatures varied resulting in groups using different cell markers to identify particular cell populations (Rezza et al., 2016).

It is also worth noting that during early hair follicle development the molecular signature of a particular cell population may change dramatically with some intermediate cellular states remaining to be elucidated (Mok et al., 2019). Another confounding issue is that hair follicle development is asynchronous: guard hair follicles are induced as early as ∼E13.5, and awl, auchene hair follicles are formed at ∼E15.5, while the zigzag hair follicle, which make up about 80 % of the adult hairs, initiates morphogenesis at ∼E17.5 (Huh et al., 2013; Schlake, 2007). Our current understanding of the timing and the machinery underlying the growth of different hair follicles remains limited. Driskell et al., demonstrated that Sox2 expression in the DC may participate in controlling the hair follicle type, as evidenced by the fact that from E18.5 SOX2 expression is confined to a subset of cells (guard/awl/auchene dermal papillae, G/AA-DP, but not zigzag dermal papillae, ZZ-DP). This provides evidence that DP cells in different type of hair follicles are heterogeneous (Driskell et al., 2009). Supporting such a notion *Sox18* ablation in mice results in the loss of zigzag hairs (James et al., 2003; Pennisi et al., 2000). Conversely, Chi et al., demonstrated that the number of DP cells in the hair follicle correlates with the hair follicle type during development, suggesting that the different number of DPs induced different cumulative signaling, which then specifies the hair size and type (Chi et al., 2013).

Tackling these problems using scRNA seq, two recent back-to-back studies have uncovered new intermediate states that form during DC specification (Gupta et al., 2019; Mok et al., 2019). This has helped fill in the gap regarding the underappreciated intermediate cellular states occurring during cell fate determination. It also has provided new molecular insights into the cellular heterogeneity within particular cell populations during hair follicle development. Here, also by utilizing the same scRNA seq technology, we generated 15,086 single cell transcriptomes from E13.5, E16.5, and P0 mouse dorsal skin. This encompasses hair follicle induction, organogenesis, and the cytodifferentiation stages. By using tSNE clustering, we identified 9 major cell populations. Based on Monocle pseudotime ordering analysis and the SCENIC regulon inferring assay, we successfully constructed the epithelium/dermal cell lineage differentiation trajectory and revealed sequential activation of key regulons involved during cell fate decisions. The intercellular communications were also inferred during hair follicle morphogenesis. Taken together, our data here also provide new insights into cell fate decisions during hair follicle *in uterus* development, and more importantly, delineates molecular information regarding the underappreciated organogenesis, cytodifferentiation stages.

## Results

### Single-cell sequencing and characterization of cellular heterogeneity during hair follicle morphogenesis

To decipher the transcriptome regulatory network and cellular fate decisions during hair follicle morphogenesis, we dissociated dorsal back skin tissue from three timepoints during hair follicle development, the induction stage (E13.5), organogenesis stage (E16.5), and the cytodifferentiation stage (P0), into single cells and performed droplet-based single cell RNA seq (Fig. 1A). We detected 19,997 genes in total for E13.5 skin cells, 19,767 genes for E16.5 skin cells and 19,145 genes for P0 skin cells (Supplementary Fig. 1A and 1B). After removing low-quality cells, we obtained 15,086 single cell transcriptome profiles from E13.5, E16.5 and P0 mouse back skin cells (4,994, 5152, and 4940 single cells, respectively).

**Figure. 1.**
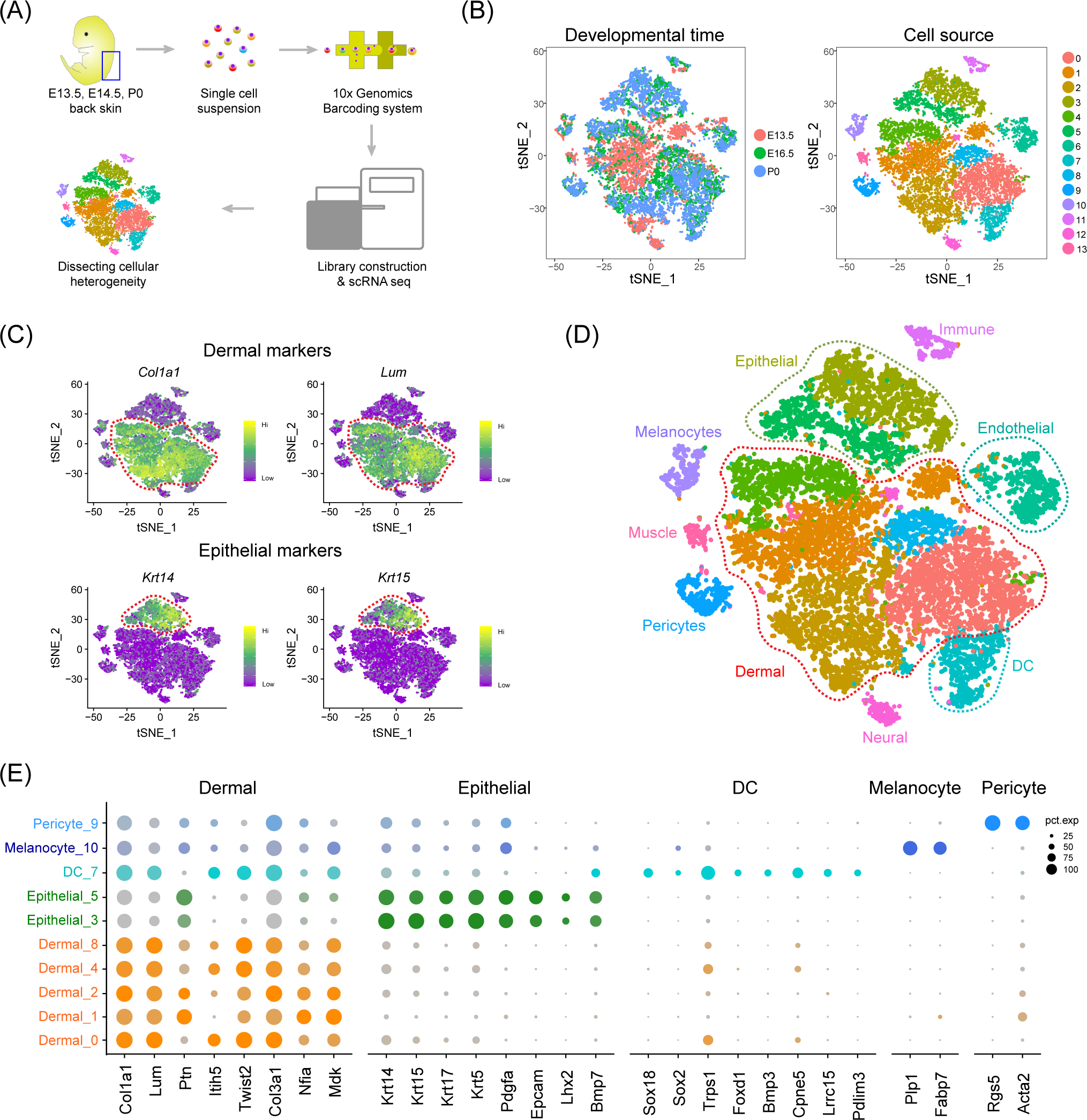
Overview of experimental procedures and characterization of major cell populations from embryonic skin tissues. (A) Schematic diagram illustrating the experimental pipeline for scRNA seq analysis of embryonic skin tissues. Single cell transcriptomes were obtained based on the 10x Chromium platform. (B) tSNE clustering of embryonic single skin cells. Each point represents one single cell and cells in the same cluster represents high similarity in transcriptome profile. The left plot depicts tSNE plot of the integrated dataset from 3 different time point and cells were color-coded with developmental time point. The right plot depicts 14 transcriptional distinct cell clusters and cells were color-coded with cluster information. (C) Visualization of dermal and epithelial marker gene expression across all single cells in the tSNE plot. *Col1a1* and *Lum* were used to mark dermal cell populations and *Krt14* and *Krt15* were used to mark epithelium populations. (D) Characterization of major cell types in the embryonic skin tissues in the tSNE plot. Cells were labeled with their cell identity and were color-coded. (E) Dot plot depicts representative dermal cell, epithelial, DC, melanocyte, and pericyte gene expression. The dot size represents the percentage of cells expressed and the color intensity represents relative expression level.

To dissect the cellular heterogeneity during hair follicle morphogenesis, we then performed tSNE clustering of all the single cells (Fig. 1B). We found that only a small percentage of the cells from E13.5 overlapped with cells from E16.5 and P0, which was consistent with the fact that dermal and epidermal cells are homogenous populations prior to differentiation (Fig. 1B, left panel) (Andl et al., 2002). Furthermore, tSNE clustering analysis revealed 14 cell clusters according to their gene expression profiles, thus preliminarily deciphering that skin cells were highly heterogeneity during hair follicle morphogenesis (Fig. 1B, right panel). To further characterize cell cluster identity, we initially performed hierarchical clustering on the 14 cell clusters (Supplementary Fig. 2A) and the results revealed 8 major branches (cluster 12; cluster 11; cluster 3, 5; cluster 10; cluster 1, 4, 8, 13; cluster 0, 2, 7; cluster 6; cluster 9). We then evaluated a series of well-recognized cell marker gene expressions and revealed 9 major cell identities: *Col1a1* and *Lum* highly expressed dermal cell clusters (Fig. 1C, top panel) (Gupta et al., 2019; Ying et al., 1997); *Krt14* and *Krt15* highly expressed epithelial cell clusters (Fig. 1C, lower panel) (Gu and Coulombe, 2007); *Pecam1* and *Kdr* highly expressed endothelial cell cluster (Detmar et al., 1998); *Sox18* and *Lef1* highly expressed DC cluster (Liu et al., 2004; Wegner and Stolt, 2005); *Plp1* and *Fabp7* highly expressed melanocyte cell cluster (Colombo et al., 2012); *Rgs5* and *Acta2* highly expressed pericyte cell cluster (Paquet-Fifield et al., 2009); *Map2* and *Stmn3* highly expressed neural cell cluster (Treutlein et al., 2016); *Myod* and *Pax7* highly expressed muscle cell cluster (Seale et al., 2000), and *Cd52* and *Fcer1g* highly expressed immune cell cluster (Supplementary Fig. 2B) (Bartling et al., 2019). These results further validated our hierarchical clustering analysis. It is of note that recent studies using scRNA on E14.5 mouse dorsal skin delineated three new DC stages (pre-DC, DC1, and DC2) (Mok et al., 2019) and our identified DC cluster specifically expressed genes here also consistent with recent observations, including the pre-DC marker *Lef1*, the DC1 marker *Prdm1*, and the DC2 marker *Inhba* (Supplementary Fig. 2B, top panel). These analyses together enable *bona fide* characterization of different cell populations within the skin tissues (Fig. 1D).

Based on Seurat analysis, we also compared cluster-specific gene expression across the cell clusters (Fig. 1E) and it was found that dermal cell clusters showed high expression levels of *Col1a1*, *Lum*, *Ptn*, *Twist2*, *Col3a1*, *Nfia* and *Mdk*, while epithelial cells showed high expression of *Krt14*, *Krt15*, *Krt17*, *Krt5*, *Pdgfa* and *Bmp7*. We also revealed DC cell cluster specific genes, including *Sox18*, *Sox2*, *Trps1*, *Foxd1*, *Bmp3* and *Cpne5*. Visualization of the top 10 expressed cluster specific genes showed obvious cluster specific expression (Supplementary Fig. 2C and Supplementary Table 1). Taken together, we have successfully identified major cell populations in the dorsal skin during hair follicle morphogenesis and identified a series of cell identity specific signature genes, which enable *bona fide* characterization of cellular heterogeneity using single cell RNA seq.

### Revealing dermal cell fate decisions during hair follicle induction stage (E13.5 – E16.5)

From E13.5 to E16.5, unspecified dermal and epidermal cells differentiate into DC and HFSC or matrix precursors, respectively (Fig. 2A). We intitially focused on the dermal cells and extracted all dermal lineage cells (dermal cell clusters and DC cluster in Fig. 1D) and performed pseudotime ordering based on the Monocle algorithm (Fig. 2B). After pseudotime analysis, we observed two branches across the hair follicle morphogenesis stage. The first branch point 2 mainly consists of cells from E16.5, while the branch point 1 consists of cells from P0 (Fig. 2B), thus deciphering major cell fate decisions during dermal cell differentiation across the hair follicle *in uterus* differentiation.

**Figure 2.**
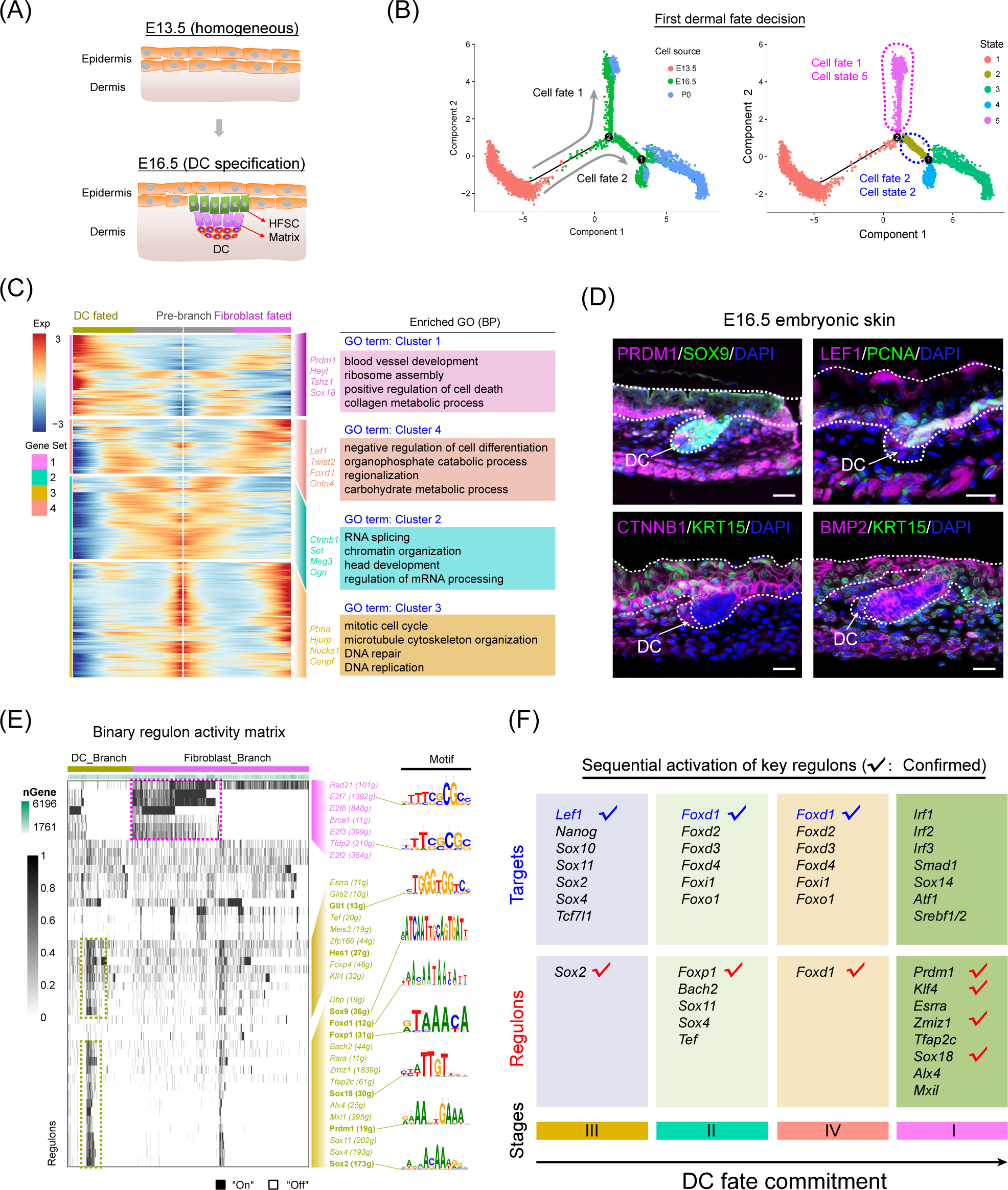
Recapitulating dermal cell fate decision towards DC fate. (A) Diagram deciphering dermal lineage and epidermal lineage differentiation at E13.5 to E16.5. (B) Pseudotime ordering of all dermal cell lineage cells. Each dot represents one cell and each branch represents one cell state. The left plot was labeled with developmental time and the right plot was labeled with cell states. (C) Heatmap illustrating the DEGs dynamics towards DC and fibroblast fate along pseudotime. The DEGs were clustered into 4 gene sets according to *k*-means and the expression curve was illustrated in the middle. GO terms enriched for each gene set were labeled in the right panel. DC fated represents cell fate 2 in Fig. 2B, while fibroblast fated represents cell fate 1 in Fig. 2B. (D) Immunofluorescence analysis of PRDM1, SOX9, LEF1, PCNA, CTNNB1, KRT15 and BMP2 expression in the E16.5 dorsal skin. Scale bars, 50 μm. (E) SCENIC binary regulon activity heatmap depicting DC and fibroblast enriched regulons. The column depicts a single cell while the row depicts regulons. For the regulons of particular interest, their representative binding motif was visualized in the right panel. “On” depicts active, while “Off” represents inactive. (F) Sequential visualization of enriched regulon activity in each gene set corresponding to Figure 2C. Their representative target genes were also provided and the red ticks depict confirmed markers in DC fate commitment.

We next focused on the branch point 2 and performed gene expression analysis along each branch (Fig. 2C). According to the pseudotime ordering analysis, dermal cells bifurcated into two cellular states (state 2 and state 5 in Fig. 2B, right panel). To reveal the sequential gene expression dynamics along each branch, we visualized gene expression dynamics along the pseudotime trajectory (Supplementary Table 2) and observed four distinct gene sets according to their expression pattern (Fig. 2C and Supplementary Fig. 3A). For the pre-branch (gene set 3), namely unspecified dermal cells from E13.5, they showed high level expression of *Ptma*, *Hjurp*, *Nucks1*, *Cenpf*, and enriched GO terms of “mitotic cell cycle, microtubule cytoskeleton organization, and DNA repair”, which was similar to the fibroblast branch. While for the DC fated branch, we found three subsequent stages (gene set 2, 4, 1). For gene set 2, we observed elevated expression of *Igfbp5*, *Set*, *Meg3* and these genes enriched GO terms of “RNA splicing and chromatin organization”, which may represent dermal cells that have received induction signals and are preparing for their subsequent differentiation. For the gene set 4, we observed high expression of *Lef1*, *Twist2*, *Foxd1*, *Cntn4* (Fig. 2C and Supplementary Fig. 3A), and these genes enriched the GO terms of “negative regulation of cell differentiation, organophosphate catabolic process, and regionalization”. It is of interest that *Lef1* and *Twist2* have been recently identified as pre-DC markers by Mok et al., and it is therefore plausible that gene set 4 represents a pre-DC stage signature gene list (Mok et al., 2019). After the pre-DC stage, our analysis revealed that DC fated cells showed a significantly high level expression of *Prdm1*, *Heyl*, *Tshz1*, *Sox18* (Fig. 2C and Supplementary Fig. 3B) and enriched the GO terms of “blood vessel development, ribosome assembly, and positive regulation of cell death”, which further emphasized their DC identity. Furthermore, when evaluating cell cycle-related gene expression during bifurcation, we also found that pro-proliferative genes, such as *Pcna*, decreased in the DC branch while the cell cycle inhibitor gene *Btg1* increased along the DC fate commitment (Supplementary Fig. 3B) (Kurki et al., 1986; Zhu et al., 2013). This is consistent with recent findings that cell cycle exit was a marker of acquisition of DC fate.

**Figure 3.**
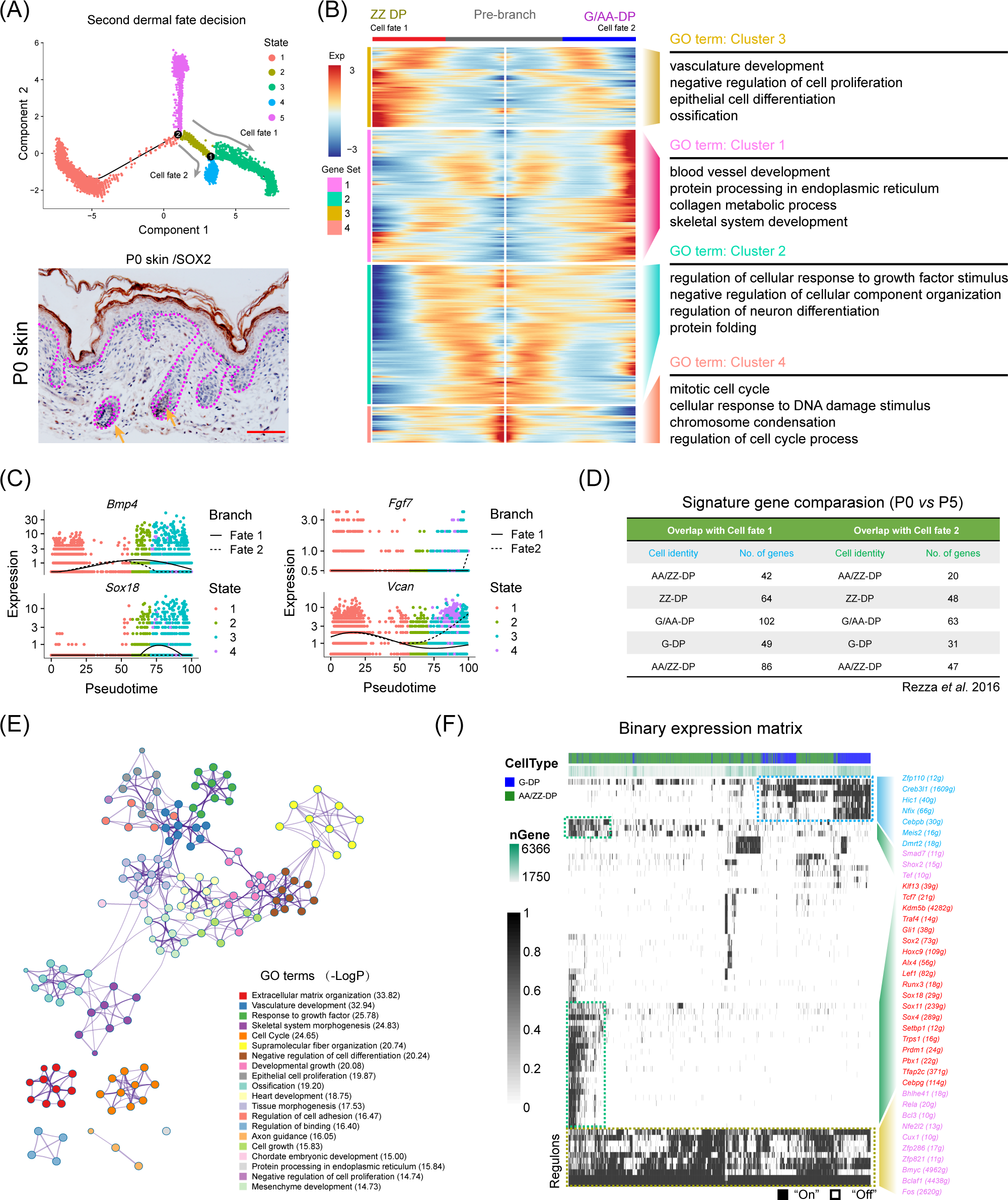
Dissecting DP population heterogeneity in the late stage of hair follicle development. (A) Monocle pseudotime trajectory construction analysis and immunohistochemistry analysis of SOX2 expression in P0 skin. Arrows indicate the SOX2 positive DP population. Scale bars, 25 μm. (B) Heatmap illustrating DP signature gene dynamics from pre branch to ZZ DP and G/AA-DP fate. The corresponding GO terms for each gene set were listed in the right panel. (C) ZZ DP makers *Bmp4*, *Sox18* expression and G/AA-DP markers *Fgf7*, *Vcan* expression along pseudotime. Cells were color-coded with cell states and the solid line represents cell fate 1, while the dashed line represents cell fate 2. (D) Comparison of DP signature genes in this study with previously identified different DP signature genes using bulk RNA seq. No. of genes depicts the number of overlapped genes. (E) GO interaction network constructed from different gene set enriched GO terms. Different colors in the network depict different GO terms. (F) SCENIC binary regulon activity heatmap deciphering G/AA-DP and ZZ DP branch specific enriched regulons. “On” depicts active, while “Off” represents inactive.

Immunofluorescence analysis also indicated that DC expressed PRDM1 and LEF1, but not PCNA, further emphasizing that DC cells exit the cell cycle at this stage (Fig. 2D). The expression of CTNNB1 and BMP2 was also detected in the DC cell populations while K15 was mainly expressed in the epidermal cells which was also consistent with our analysis here. Taken together, our analysis here successfully recapitulated the *bona fide* dermal cell fate decision to a DC fate, and revealed the transcriptome landscape during DC fate commitment.

After recapitulation of the dermal cell transcriptome landscape during the hair follicle induction stage, we then investigated the key transcriptional factors involved during the first dermal cell fate decision. We implemented a single-cell regulatory network inference and clustering (SCENIC) pipeline to infer key regulons involved during the hair follicle dermal cell fate decision (Aibar et al., 2017). The SCENIC algorithm revealed a series of key regulons and their corresponding target genes (Supplementary Table 3). For the DC branch cells, they enriched regulons such as *Gli1*, *Hes1*, *Sox9*, *Foxd1*, *Foxp1*, *Sox18*, *Prdm1*, and *Sox2* (Fig. 2E), all of which were also defined markers during DC fate commitment (Mok et al., 2019). Our immunofluorescence analysis also verified the expression of SOX9 and SOX18 in the DC population (Fig. 2D and Supplementary Fig. 3C). We also revealed other candidate regulons which may play vital roles during DC specification, such as *Glis2*, *Zfp160*, *Zmiz1,* and *Sox11,* etc. Combining our Monocle cell fate comparison assay with our regulon inferring assay, we further analyzed sequential activation of the key regulons (Fig. 2F), and it was revealed that *Sox2*, *Foxp1*, *Foxd1* were early activated regulons, targeting to *Lef1*, *Foxd* family members, and *Foxo1*. Noteworthy, these regulons illustrated above and their corresponding targets have been recently identified as preDC signature genes (tick labeled), further demonstrating their roles during the DC fate commitment.

### Psedotime ordering analysis reveals that G/AA-DP and ZZ DP are transcriptome distinct branches

Since we successfully recapitulated the dermal cell lineage pseudotime differentiation trajectory and delineated DC cell fate commitment prior to E16.5, we then focused on the next development trajectory from E16.5 to P0 (Fig. 3A, top panel). At which time both DP cells are specified, including guard hair follicles, Awl/Auchene hair follicles, and Zigzag hair follicles (Huh et al., 2013; Tsai et al., 2014). Interestingly, consistent with Driskell et al., our immunohistochemistry analysis also showed that SOX2 was expressed in the G/AA-DP, but not the ZZ-DP (Fig.3A, lower panel). Pseudotime ordering analysis revealed that DP cells bifurcated into two branches, also suggesting that two distinct DP populations exist in P0 skin hair follicles (Fig. 3A, top panel). To further identify signature genes between the different DP cell populations, we used Monocle to perform differential gene expression (DEG) analysis between the two branches (Fig. 3B). Our analysis identified 606 signature genes for cell fate I and 1,004 signature genes for cell fate 2. Specifically, we found that branch 1 expressed high levels of *Bmp4*, *Sox18*, *Fgfr1*, *Gli1*, *Lef1*, and *Notch1*, while branch 2 showed high expression of *Fgf7*, *Vcan*, *Wnt9a*, *Notch1*, *Dlk1*, *S100b*, and *Sod3* (Fig. 3C and Supplementary Fig. 4A). For cell fate 1, DEGs enriched the GO terms of “vasculature development, negative regulation of cell proliferation, and epithelial cell differentiation”, while for cell fate 2, DEGs enriched the GO terms of “blood vessel development, protein processing in the endoplasmic reticulum, and collagen metabolic process”. Several groups demonstrated that *Sox18* specifically expressed in zigzag hair follicle DP (Graham et al., 2003; Taranova et al., 2006), and ablation of *Sox18* has been demonstrated to reduce zigzag hair formation (James et al., 2003; Pennisi et al., 2000). Therefore, we termed cell fate 1 as the ZZ-DP fate (Fig. 3B and Supplementary Fig. 4B, left panel). For cell fate 2, Driskell *et al*. demonstrated that *Fgf7* significantly increased in G/AA-DP compared with ZZ-DP (Driskell et al., 2009), we therefore termed cell fate 2 as the G/AA-DP fate (Fig. 3B and Supplementary Fig. 4B, right panel).

We then compared previously reported DP signature genes with our Monocle analysis identified G/AA-DP and ZZ-DP signature genes. As far as we know, little transcriptome information is available for P0 DP from different hair follicles, we therefore compared our identified DEGs with previously identified DP signature genes from P5 hair follicles (Fig. 3D and Supplementary Table 4). Unexpectedly, it was found that DEGs from ZZ-DP or G/AA-DP overlapped similarly with previously identified 5 DP populations. However, it’s notable that the branch endpoint showed similar expression of *Sox18*, *Bmp4*, and *Vcan*, while the differences mainly concentrated intermediate cells en route to the end branch (Supplementary Fig. 4B). It is therefore plausible that intermediate states may exist during the underappreciated DP specification stage *prior to* entering the hair cycle, which may also account for the inconsistent expression reported by different groups. We further visualized several core DP markers and also observed a similar expression pattern, including *Fgf7*, *Lef1*, *Gli1*, and *Notch1* (Supplementary Fig. 4C) (Rezza et al., 2016).

To further capture the relationships between the terms enriched from the different gene sets, we then visualized the enrichment network (Fig. 3E) and compared their shared GO terms (Supplementary Fig. 4D). It was found that the four gene sets shared many co-enriched GO terms, and the top enriched GO terms included extracellular matrix organization, vasculature development and response to growth factors. We also applied SCENIC to infer key regulons involved during the DP cell fate decision (Fig. 3F). It was found that both DP populations enriched regulons such as *Bmyc*, *Bclaf1*, and *Fos*, while the ZZ branch specifically enriched regulons such as *Gli1*, *Sox2*, *Lef1*, *Sox18,* and *Prdm1*, and the G/AA-DP branch enriched regulons such as *Zfp110*, *Creb3l1*, *Meis2*, and *Dmrt2*. The differences found between regulon enrichment also suggests the heterogeneity of the ZZ-DP and G/AA-DP.

### Recapitulating epithelial cell fate decisions towards matrix and HFSCs precursors (E13.5 – E16.5)

After delineating the dermal cell fate decisions, we then investigated the underappreciated epidermal cell lineage fate decisions. Histology analysis showed that E16.5 mouse dorsal skin had obvious primary follicle structures (Fig. 4A). We then extracted epithelium cells from Seurat and performed Monocle cell trajectory analysis using variable genes identified by Seurat as ordering genes (Fig. 4B, left and Supplementary Fig. 5A, 5B). Cell trajectory analysis revealed that epithelium cells also showed two main bifurcation points along the cell trajectory. The first bifurcation point consists of cells derived from E16.5, while the second bifurcation point mainly consists of cells derived from newborn mouse dorsal skin (Fig. 4B, right).

**Figure 4.**
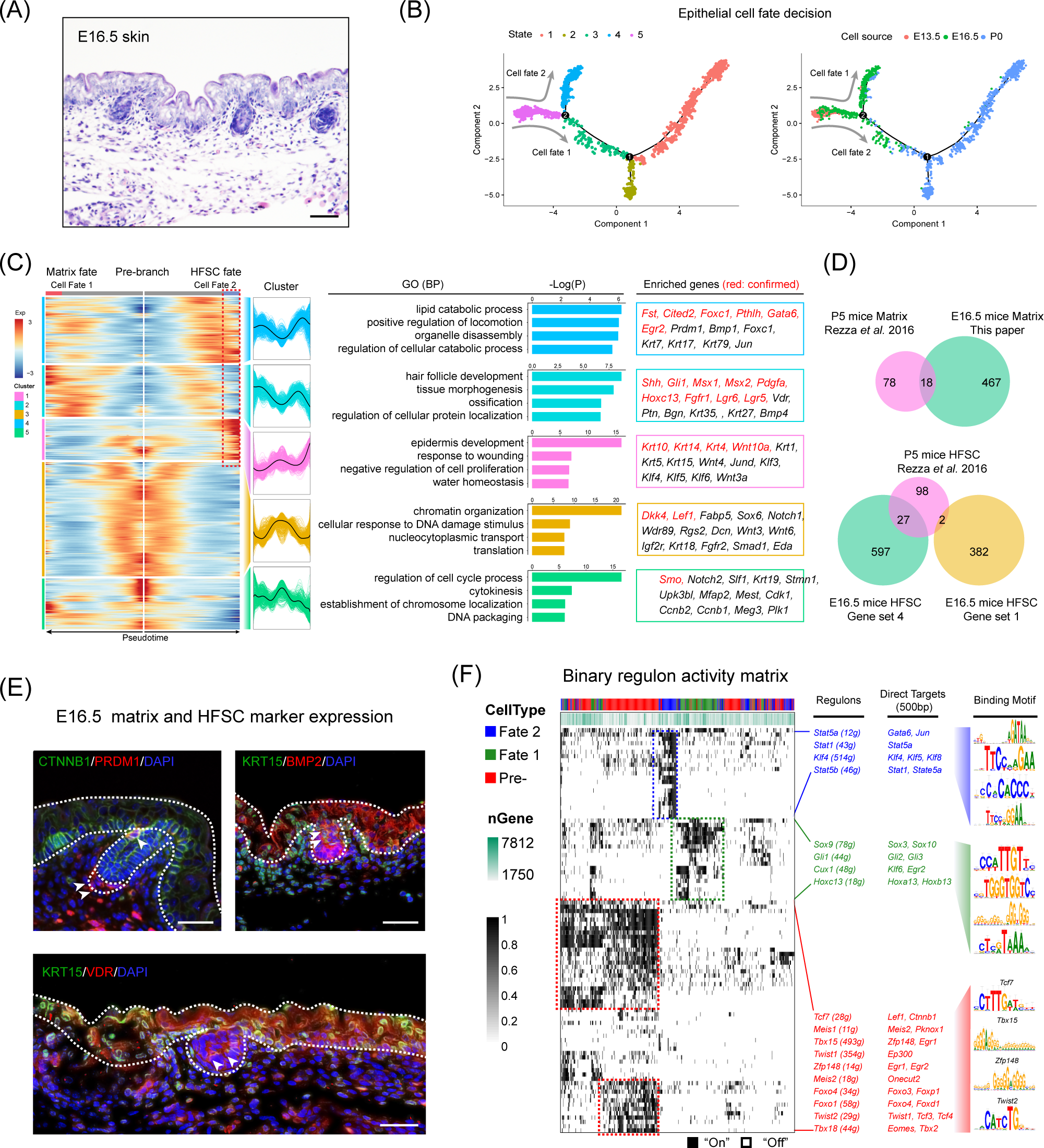
Dissecting molecular signature underlying matrix and HFSC precursor fate commitment. (A) Histology analysis of E16.5 embryonic skin. Scale bar, 50 μm. (B) Pseudotime ordering of all epithelium cell populations from three developmental time stages. Each dot represents one cell. The left plot was color-coded with stage information, while the right plot was color-coded with developmental stages. (C) Heatmap displaying branch specific DEGs expression in branch point 2 in Figure 4B. Cell fate 1 indicates matrix fate and cell fate 2 indicates HFSC fate. The corresponding expression curve and enriched GO terms for each gene set were visualized in the middle panel. The representative DEGs for each gene set were shown in the rectangular box (right panel) with red depicting confirmed signature genes. (D) Matrix and HFSC precursor signature gene comparison between this study and Rezza et al., using bulk RNA seq for P5 dorsal skin tissues. (E) Immunofluorescence analysis of CTNNB1, PRDM1, KRT15, BMP2 and VDR expression in E16.5 skin tissues. Scale bar, 50 μm. (F) Binary regulon activity heatmap illustrating the branch-specific enrichment of key regulons. The representative regulons and their corresponding targets (500 bp upstream of TSS) were listed in the middle panel. The binding motif was listed in the right panel.

We initially focused on the first bifurcation point and performed DEG analysis comparing each branch (cell fate 1 *vs* cell fate 2 in Fig. 4B). DEG analysis revealed the two branches as matrix progenitor cells and HFSC progenitor cells as evidenced by the high expression of canonical markers (Fig. 4C and Supplementary Fig. 5C). For the HFSC branch, the DEGs enriched two distinct gene clusters (cluster 1 and 4) (Fig. 4C, indicated the red rectangular box). For cluster 4 they enriched canonical markers such as *Fst*, *Cited2*, *Foxc1*, *Sox9*, *Gata6* (Supplementary Fig. 5C) (Greco et al., 2009; Nowak et al., 2008), and enriched the GO terms of “lipid catabolic process, positive regulation of locomotion, and organelle disassembly”. Conversely for gene cluster 1 they enriched genes such as *Krt10*, *Krt14*, *Krt4*, *Wnt10a* and the GO terms “epidermis development, response to wounding, and negative regulation of cell proliferation”. For the matrix cell populations our analysis also delineated classic matrix progenitor markers, including *Shh*, *Hoxc13*, *Msx1*, *Msx2*, *Lgr5*, *Lgr6*, *Pdgfra* (Supplementary Fig. 5C) (Greco et al., 2009; Panteleyev et al., 2001; Rezza et al., 2016) with, *Vdr*, *Lhx2*, *Ptn*, *Klf5* and *Bmp4* also differently expressed in the matrix progenitors. GO enrichment demonstrated that matrix progenitors enriched the GO terms “hair follicle development, tissue morphogenesis, and ossification.” We then compared our enriched DEGs with matrix and HFSC progenitor signature genes identified by Rezza et al., (E16.5 *vs* P5) (Fig. 4D and Supplementary Table 5). It was found that for the matrix cell populations about 18.75 % DEGs co-expressed with our analysis. Interestingly, for the HFSC population we found that 21.6 % DEGs identified by Rezza et al. overlapped with gene set 4 (Fig. 4D and Supplementary Table 5), while only two genes overlapped with gene set 1. This demonstrated that gene set 1 may indicate an underappreciated HFSC state. Immunofluorescence analysis also confirmed the expression of PRDM1, BMP2, KRT15 in the HFSC population and VDR in the matrix population (Fig. 4E).

Next, we performed SCENIC regulon inferring analysis and obtained a list of candidate cell state specific regulons involved in the matrix progenitor and HFSC progenitor cell specification (Fig. 4F). For the unspecified epidermis (state 5) our DEG analysis showed that *Dkk4* and *Lef1* showed high expression levels in the pre-branch gene set (Supplementary Fig. 5D), which consisted of all epithelial cells derived from the E13.5 dorsal skin. SCENIC analysis indicated that regulons such as *Tcf7*, *Tbx15*, *Foxo1*, *Foxo4* were enriched in the pre-branch gene set. Noteworthy, *Lef1* and *Ctnnb1* (β*-catenin*) were two direct targets of *Tcf7* and together with the fact that *Ctnnb1* knockout mice fail to form DC during hair follicle development (Tsai et al., 2014), it is plausible that *Tcf7* acts in concert with downstream *Lef1* and *Ctnnb1* and may be a regulator controlling DC formation. We also delineated a series of other regulons and their corresponding targets including *Tbx15* (targets to *Zfp148*, *Egr1*), *Zfp148* (targets to *Egr1*, *Egr2*), Foxo signaling members *Foxo4* and *Foxo1* and twist signaling members *Twist 1, 2* (target to *Tcf 3/4*, *Ep300*), all of which have been described in previous research. For the matrix progenitor branch our analysis demonstrated that *Sox9*, *Hoxc13*, *Gli1,* and *Cux1* were core regulons involved, while for the HFSCs fate we delineated that *Stat* family members *Stat1*, *Stat5a*, *Stat5b* and Krüppel-like factor gene family members *Klf4*, *Klf5*, *Klf8* were core identified regulons.

### Revealing hair shaft and IRS fate decisions during cytodifferentiation (E16.5 – P0)

After recapitulating the key events involved at the hair follicle organogenesis stage (around E16.5), we then focused on the next cytodifferentiation stage (around P0) (Fig. 5A). As expected, single cell trajectory analysis of the P0 epithelium cells bifurcated into two branches (Fig. 5B). To infer their cell identity, we similarly performed DEG analysis between the branches using Monocle. It was found that cell fate 2 significantly enriched IRS related marker genes including, *Gata3*, *Notch1*, *Krt16*, *Wnt7b* and *Scube2*, while cell fate 1 enriched hair shaft related genes, including *Lhx2*, *Shh*, *Hoxc13*, *Mycl*, *Myb*, *Nrp2*, *Casz1*, and *Edar* (Supplementary Fig. 6A) (Yang et al., 2017). These data together demonstrated that we have successfully recapitulated the hair shaft and IRS differentiation trajectory from the matrix progenitors. To further unmask the gene expression profile and gene regulatory network underlying hair shaft and IRS specification, we performed DEG analysis and SCENIC regulon inferring analysis along with the Monocle trajectory analysis (Fig. 5C, 5D). Monocle branch specific gene expression analysis revealed hair shaft enriched genes such as *Shh*, *Hoxc13*, *Msx1/2,* and *Bmp4*, all of which had been well characterized to play vital roles during hair shaft differentiation (Millar, 2002; Yang et al., 2017). We also found a series of other DEGs such as *Krt25*, *Krt71*, *Mycl*, *Myb*, *Lhx2*. We further compared our Monocle identified DEGs with Anagen II hair shaft/IRS signature genes identified by Yang et al. (Supplementary Fig. 6B). We identified 585 branch specific DEGs for hair shaft and about 3.76% (22/585 genes) overlapped with the anagen II hair shaft signature genes, thus it is plausible that the hair shaft cells display distinct gene expression patterns prior to entering the hair cycle (Supplementary Table 6). Similarly, IRS gene set 1 shared 2.75% (10/363) and IRS gene set 3 shared 3.23% (16/495) overlapping genes with the anagen IRS signature genes. These data together emphasized that embryonic hair shaft and the IRS shared distinct gene expression profiles compared with the anagen hair shaft and the IRS.

**Figure 5.**
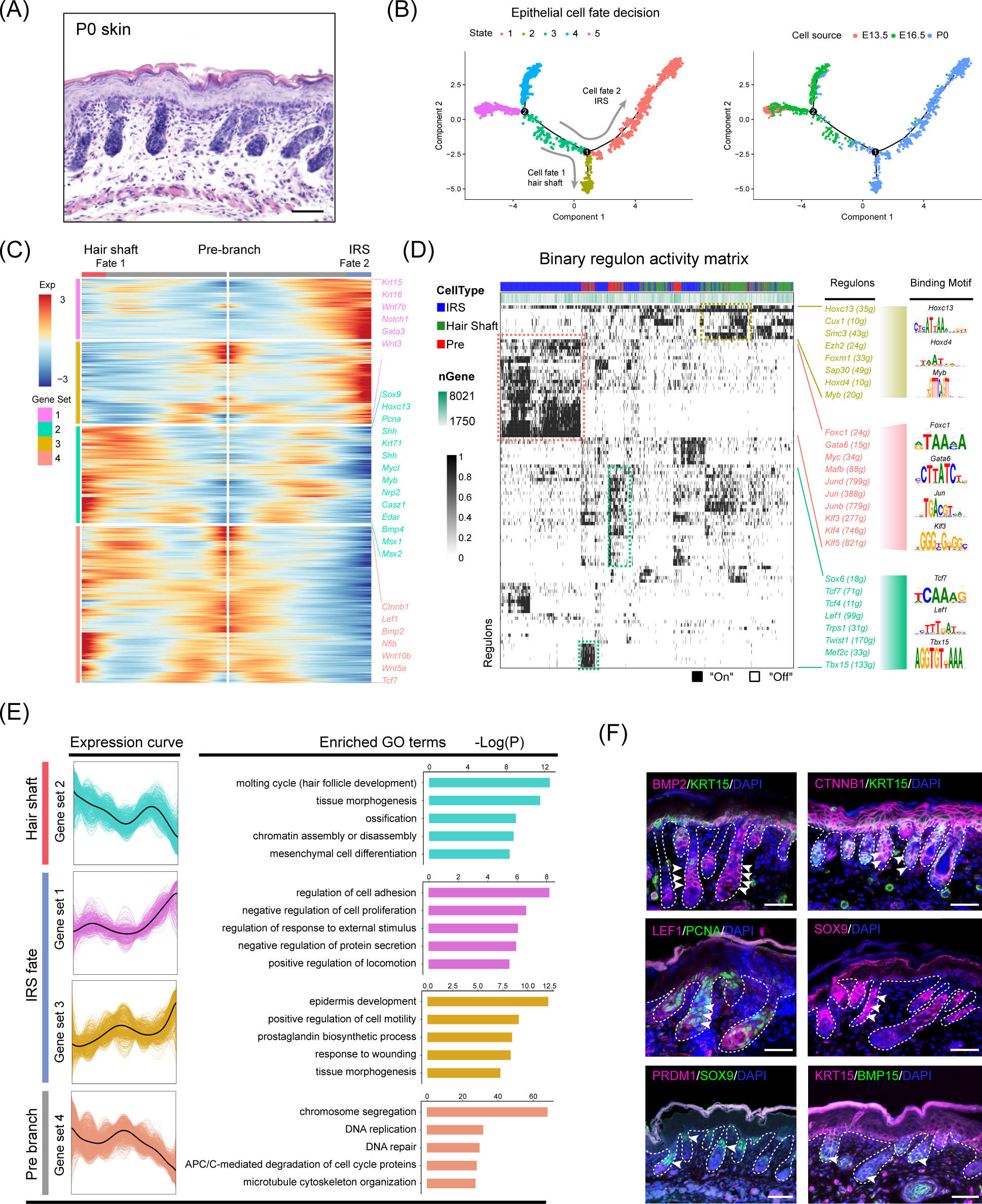
Dissecting hair shaft and IRS fate commitment from matrix precursors. (A) Histology analysis of P0 mouse skin. Scale bar, 50 μm. (B) Pseudotime visualization of the hair shaft and IRS fate decisions in the epithelium single cell pseudotime trajectory. (C) Heatmap illustrating branch specific DEGs dynamics along pseudotime. Cell fate 1 indicates hair shaft fate and cell fate 2 indicates IRS fate. (D) Binary regulon activity heatmap demonstrating branch specific enrichment of regulons. The cell states correspond to Figure 5B. The representative regulon and binding motifs were listed in the right panel. (E) DEGs expression dynamics and regulon enrichment in each gene set. The gene expression curve and GO terms for each gene set were listed in the left panel, the gene set enrichment of regulons and their corresponding targets were listed in the right panel. Red indicates confirmed signature genes. (F) Immunofluorescence analysis of hair shaft markers BMP2, CTNNB1, LEF1, SOX9, IRS markers KRT15, and PCNA, BMP15 in P0 skin. Scale bar, 25 μm.

We then used SCENIC to infer transcriptional factor (TF) regulatory information among the three branches (Fig. 5D). SCENIC regulon inferring analysis revealed that the hair shaft branch (State 2) enriched regulons such as *Hoxc13*, *Cux1*, *Hoxd4*, and *Myb*. The expression of *Hoxc13* had been long demonstrated as a key transcriptional factor in promoting hair shaft specification. Besides, our analysis here also unmasked a series of significantly enriched regulons for the underappreciated IRS (State 1) which included *Gata6*, *Foxc1*, *Jun* family members (*Jun*, *Junb*, *Jund*), and *Klf* family members (*Klf3/4/5*). Noteworthy, *Gata6* has been previously demonstrated as a marker for the IRS and perturbation of *Gata6* caused dilation of the hair follicle canal demonstrating an indispensable role during hair follicle morphogenesis (Swanson et al., 2019).

To gain further insight into the gene regulatory machinery underlying the matrix progenitors’ commitment to the hair shaft or IRS, we then compared their gene expression profiles along the pseudotime trajectory (Fig. 5E). We first compartmentalized the identified branch specific DEGs using k-means clustering. The hair shaft significantly enriched gene clusters (cluster 2) enriched GO terms of “molting cycle (hair follicle development), tissue morphogenesis, and ossification”. Previously, Yang et al. demonstrated that signature genes of anagen II hair shaft had the enriched GO terms of “cholesterol biosynthetic process, steroid biosynthetic process and lipid metabolic process, which further emphasized that the hair shaft showed distinct gene expression patterns *prior to* entering the hair cycle. Similarly, the IRS highly expressed genes (cluster 1, 3) had the enriched GO terms “regulation of cell adhesion, negative regulation of cell proliferation (cluster 1) and epidermis development, positive regulation of cell motility, and prostaglandin biosynthetic process,” which were also different from Yang et al. which had the anagen IRS signature genes enriched GO terms “DNA replication, cell cycle, and multicellular organism development. Last, to confirm our analysis, we performed immunofluorescence showing that BMP2, CTNNB1, LEF1, and SOX9 expressed in the hair shaft, while KRT15 highly expressed in the IRS (Fig. 5F). Furthermore, we found that hair shaft showed a high expression of the cell proliferation marker PCNA while BMP15 was mainly expressed in the DP cells, while PRDM1 was not detectable in hair follicles at P0. Taken together, our analysis here provides underappreciated information during the matrix cells fate commitment to the hair shaft and IRS.

### Ligand-receptor interaction prediction during hair follicle morphogenesis

Since we have successfully recapitulated dermal and epidermal cell fate decisions and delineated the molecular signatures of the different cell populations, we then used a public ligand-receptor database to infer intercellular communications during early hair follicle development (Ramilowski et al., 2016; Skelly et al., 2018). By comparing the cell identity specific genes with ligand-receptors, we sorted hypothetical ligand-receptor pairs among different cell populations (Fig. 6). For E13.5 to E16.5 ligand-receptor pairs (DC specification and epidermal specification to the matrix and HFSC), we found stronger interaction relationships among matrix, HFSC, and DC populations at E16.5 (Fig. 6A). Noteworthy, we observed robust ligand-receptor pairs within the DC population including, *Vcan*, *Egfr*, and *Bmp7*, indicating a strong autocrine relationship at this stage. In E13.5 dermal and epidermal cell populations we also observed strong intercellular communication. Specifically, the ligand *Itgb1* was expressed in the dermal population while its ligands were mainly expressed in the epidermal cells.

**Figure 6.**
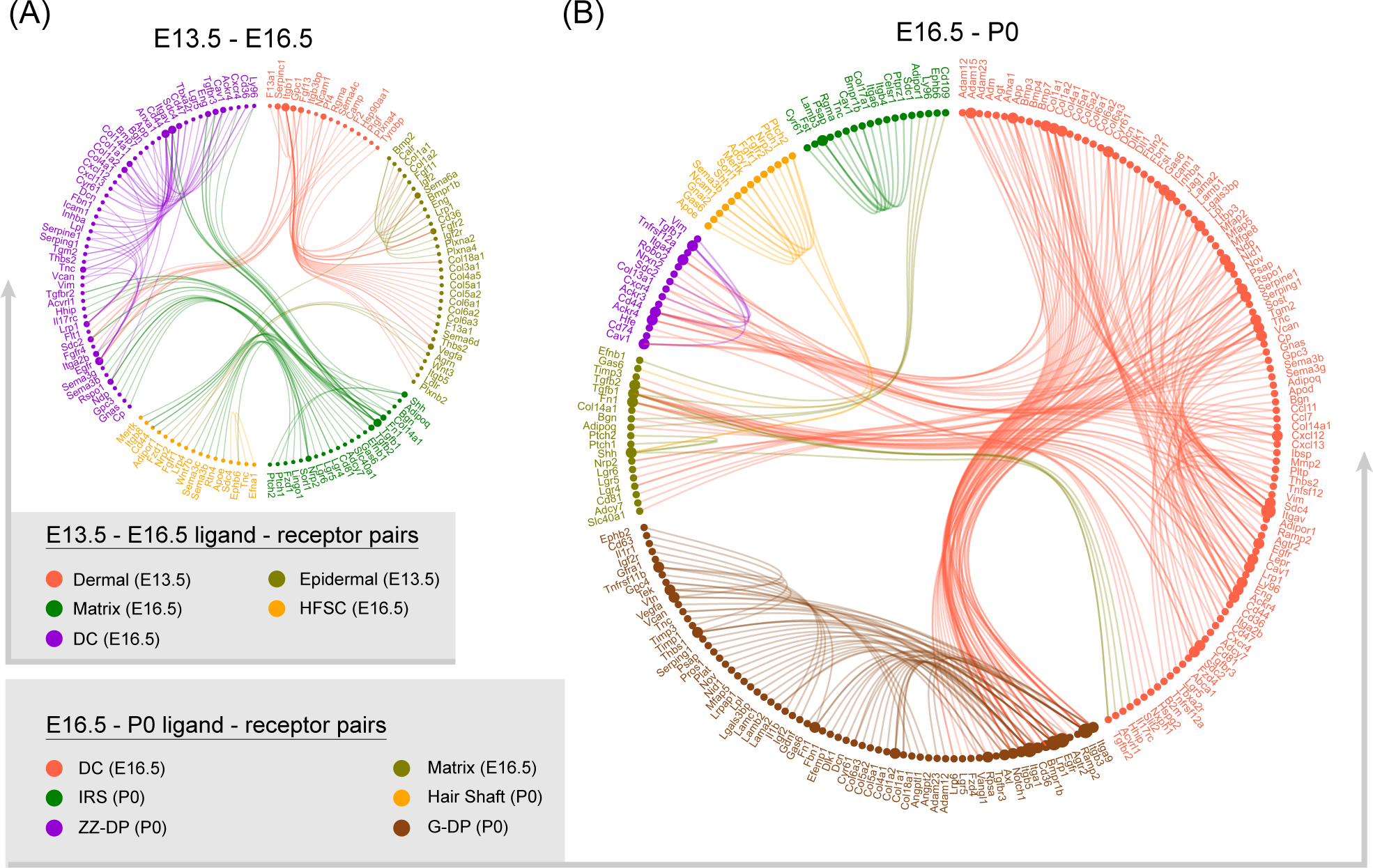
Intercellular ligand-receptor prediction. Ligand-receptor pairs between E13.5-E16.5 (A) and E16.5-P0 (B) main cell populations. Different cell populations were color-coded and ligand-receptor pairs were linked with a solid line.

At the hair follicle cytodifferentiation stage, we inferred potential ligand-receptor pairs using priori knowledge. Our analysis showed strong intercellular communication between E16.5 DCs and P0 G-DP cells including the BMP signaling members *BMP3*, *4*, and *7*, Notch signaling ligands *Jag1*, *Dlk1,* and collagens family members *Col1a1*, *Col1a2*, *Col4a1*. This further emphasized the indispensable roles of these well-defined pathways during DP specification. We also observed strong intercellular communication between E16.5 DC and P0 ZZ-DP cells. However, some of these ligand-receptor pairs differ between E16.5 DC and P0 G-DP, further suggesting heterogeneity in DP cells as we illustrated in our DEG analysis. For IRS, hair shaft and G-DP, we also observed strong autocrine signaling as revealed by abundant ligand-receptor pairs within the same cell population. These analyses together showed that strong intercellular communication was involved during early hair follicle development.

## Discussion

The lack of bona fide markers characterizing key cell populations and the asynchronous development of different hair follicles during early development has produced two major obstacles hindering our current understanding of hair follicle development. However, seminal works have revealed the molecular signatures of different hair follicle cells and their roles during hair follicle development. Although, the conclusion raised by different groups varies, two major questions remain to be answered: First, the *bona fide* characterization of molecular signatures of key cell populations involved during hair follicle morphogenesis, and second, the level of heterogeneity within DP populations, which may be responsible for the asynchronous growth of the hair follicles.

With the development of high-throughput scRNA-seq analysis, many complicated biological processes have been delineated in an unprecedented manner. This has been particularly beneficial in the area of organogenesis research as scRNA seq has the robust ability to deconstruct cell heterogeneity within complicated tissues (Kamimoto and Morris, 2018). Very recently, by using scRNA seq two back-to-back papers published in *Developmental Cell* have delineated an underappreciated intermediate pre-DC fate transition stage occurring prior to DC formation (induction stage, E13.5 - E14.5) (Gupta et al., 2019; Mok et al., 2019). Here, utilizing the same technology we performed scRNA seq on hair follicles at three-time points that encompass all three stages during hair follicle development (induction, organogenesis, and cytodifferentiation) and provide comprehensive knowledge on the molecular events involved. For DC fate commitment in our analysis, we observed two different gene sets en route to a DC fate (Fig. 2C, gene set 1 and 4). Interestingly, *Lef1* and *Twist2* were enriched in gene set 4, while *Prdm1* and *Sox18* were enriched in gene set 1. This was consistent with Mok *et al*., with transcription factors *Lef1* and *Twist* expressed in the pre-DC stage (Mok et al., 2019) while transcription factors *Prdm1* and *Sox18* expressed in the DC1 stage, thus deciphering the bona fide characterization of an intermediate DC stage. We also identified other signature genes including *Tshz1* (a transcriptional regulation of developmental processes), *Heyl* (an effector of Notch signaling and a regulator of cell fate decisions), and *Cntn4* (a glycosylphosphatidylinositol-anchored neuronal membrane protein) which may function as new markers for DC cell populations. Our analysis here is also consistent with recent findings that DC fate commitment requires cell cycle exit as evidenced by upregulated expression of the cell cycle inhibitor *Btg1* and downregulated expression of the pro-proliferative genes *Pcna* (Biggs et al., 2018). These together further emphasize the presence of intermediate DC stages and that cell cycle exit is one of the marker events during DC specification.

We also delineated the epithelium fate commitment to HFSC and matrix precursors which to our knowledge has not been comprehensively reported (Millar, 2002). By using Monocle pseudotime ordering analysis, we successfully recapitulated the HFSC and matrix cell differentiation trajectory and found that matrix cells at this stage-enriched genes such as *Shh*, *Pdgfra*, *Gli1*, *Hoxc13*. Interestingly, the expression of *Shh*, *Pdgfra* in matrix cells has been demonstrated to promote down growth via *Gli1*, *Gli2* (Sennett and Rendl, 2012) which was consistent with our analysis here. Our GO analysis revealed that “hair follicle development, tissues morphogenesis” were enriched in matrix signature genes, further deciphering the molecular events during the envelopment of the DC cells by the matrix cell populations. Interestingly, the expression of *Hoxc13*, a key component involved in hair shaft differentiation, had been detected as early as E16.5 indicating an earlier activation of hair shaft fate than previously understood. It would be interesting to investigate whether these intermediate states exist during hair shaft fate commitment in future studies. Our analysis showed that HFSC fate commitment involved canonical HFSC markers such as *Fst*, *Cited2*, *Foxc1*, *Pthlh*, and *Gata6*. DEG analysis revealed 2 gene sets similar to the DC fate commitment which were reminiscent of the intermediate preDC stage prior to DC fate commitment. For gene set 2, DEGs enriched the GO terms “lipid catabolic process and positive regulation of locomotion”, while gene set 1 enriched the GO terms “epidermis development and response to wounding”, displaying distinct molecular pathways are involved. Therefore it is plausible that the intermediate cellular stages may exist during HFSC fate commitment and future studies may focus on this topic.

As development continues matrix cells give rise to the hair shaft and IRS populations, which was consistent with our Monocle analysis here (Fig. 5B). There is limited information regarding the molecular signatures of key cell populations in the late two stages of hair follicle development. Here we successfully delineated hair shaft and IRS signature gene expression profiles and the gene functional categories involved during their fate commitment from matrix precursors. For hair shaft fate commitment our analysis showed that *Hoxc13*, *Shh*, *Bmp4*, *Msx1/2* were specifically enriched, which was consistent with previous findings (Millar, 2002). However, the expression of *Notch1* and *Lef1* was not enriched in cells with a hair shaft fate but in cells with the IRS fate and the pre branch fate, respectively. For GO enrichment, hair shaft had enriched GO terms involved in molting cycle, hair cycle, and hair follicle development. This showed distinct differences compared with the anagen hair shaft, thus emphasizing distinct gene expression profiles after entering the hair cycle. Interestingly, for IRS fate specification, our analysis also observed two gene sets. Gene set 1 enriched the GO terms “regulation of cell adhesion and regulation of cell-cell adhesion” and gene set 3 enriched the GO terms “epidermis development and skin development”. This may also indicate different intermediate stages prior to IRS fate commitment. Taken together, our data here provides an unprecedented insight into the gene expression profiles of hair shaft and IRS cell populations during the late stages of hair follicle development. More importantly, the heterogeneity within these particular cell populations was clearly revealed in our research and may also account for the discrepancy in gene expression signatures previously reported.

Unexpectedly, pseudotime ordering of all dermal cell lineages revealed two branch points particularly for the late stage (E16.5 - P0) which was reminiscent of the discussion of DP heterogeneity. In 2009, Driskell et al. showed that different signaling is involved during hair follicle type determination (Driskell et al., 2009), while subsequent research by Chi et al. demonstrated that the number of DPs dictated the size and shape of the hair (Chi et al., 2013). Further enhancing the latter hypothesis, Rezza et al., isolated different DP populations from P5 mice back skin based on a flow cytometery analysis and performed bulk RNA seq on different DP populations (Rezza et al., 2016). By comparing their expression profiles they demonstrated high similarities for all DP populations. However, our analysis here indicated that DP cell populations bifurcated into two distinct fates indicating DP heterogeneity. Noteworthy, for the discrepancy between reports by Driskell and Chi, it should be mentioned that Chi et al. used adult mice as a research model in which hair follicles had entered the hair cycle, which may be different from an *in uter*o situation.

Supporting such a hypothesis, our data here indicated that the IRS and hair shaft showed distinct gene expression patterns compared to those that have entered the hair cycle. Interestingly, by in-depth analyzing the DEGs between the two DP branches we found that the endpoint in each branch shared a higher similarity while the intermediate cells en route displayed high heterogeneity. It’s therefore plausible that underappreciated intermediate stages may exist during DP development. Because of this DPs showed similar molecular profiles in P5 as reported by Rezza et al. while displaying higher heterogeneity in DPs as reported by Driskell et al., in embryonic skin. It will be interesting for future studies to focus on this topic which may provide new insights into hair follicle development.

In summary, our research highlights the characterization of the underappreciated molecular signatures of embryonic hair follicle progenitors using unbiased scRNA seq. Based on single-cell transcriptome analysis we delineated key events underlying dermal and epithelium fate decisions during hair follicle morphogenesis. Our data here also provides new insights into the heterogeneity of the DP cell populations and intercellular communication during hair follicle development. These data together enable an in-depth understanding of the molecular machinery underlying embryonic hair follicle development and may also provide valuable information regarding the etiology of skin-related diseases, such as melanoma.

## Methods

### Experimental Animals

About 7-8 weeks old C57/BL6 strain mice were used in this study, all mice used were purchased from Beijing Vital River Laboratory Animal Technology Co., Ltd and were housed in a temperature-controlled room with *ad libitum* food and water. To obtain pregnant mice, female mice were mated with male mice overnight at a ratio of 3:1, and mice with the presence of vaginal plug the next morning were denoted as 0.5 day post partum (dpp). To obtain E13.5 and E16.5 fetus dorsal skin, the pregnant mice were sacrificed by cervical dislocation and the gender was determined according to the fetus ovary/testis. Male mice were used to avoid potential hair cycle variation as previously indicated (Yang et al., 2017). All experimental procedures involved in this study were approved by the Experimental Animal Manage Committee of Northwest A & F University.

### Histological analysis and Immunofluorescence Staining

The skin tissues isolated from the fetal dorsal skin were fixed with 4 % paraformaldehyde (Sorlabio, Beijing, China) at 4°C overnight. The next morning, the fixed tissues were then dehydrated in an ethanol solution and further incubated with xylene for 30 min. After incubation with xylene the samples were embedded in paraffin blocks. The embedded paraffin blocks were cut with a Leica RM2255 microtome (Leica, Nussloch, Germany) at a thickness of 5-7 μm and the samples were transferred to APES (ZSGB-BIO, Bejing, China) treated slides to avoid detachment.

For hematoxylin and eosin (H & E) staining, the slides were deparaffinized in 100% xylene solutions for 30 min and further rehydrated in an ethanol series. After rehydration the slides were stained with hematoxylin solution for 7 min followed by washing twice with distilled water for 5 min. After rinsed with 1 % HCl (v/v) ethanol solution for 3 - 5 s, the slides were immediately washed with 45 °C water for 5 min. Followed by a dehydrated procedure, the slides were then stained with 1 % eosin ethanol solution and further rinsed with absolute ethanol solution for 10 min. Finally, the slides were mounted with neutral resins mounting medium and pictures were taken under an optical microscope.

For immunofluorescence staining analysis, the slides were deparaffinized in 100 % xylene solutions for 30 min and then hydrated with an ethanol series. To perform antigen retrieval, slides were incubated in boiled 0.01 M sodium citrate buffer (pH = 6.0) for 10 min and then cooled down to room temperature. Blocking was performed with 3 % BSA and 10 % donkey serum in 0.5 M Tris-HCI buffer for 30 min at room temperature, and slides were then incubated with primary antibodies at 4 °C overnight.

**Table 1.** Demographic features of the patients

|  |  |  |
| --- | --- | --- |
| Rabbit monoclonal anti-LEF1 | Cell Signaling | Cat# 2286, RRID:AB_659971 |
| Rabbit polyclonal anti-CTNNB1 | Proteintech | Cat# 51067-2-AP, RRID:AB_2086128 |
| Rabbit anti-VDR | Proteintech | Cat# 14526-1-AP, RRID:NA |
| Rabbit polyclonal anti-SOX2 | Proteintech | Cat# 11064-1-AP, RRID:AB_2195801 |
| Mouse monoclonal anti-K15 | Abcam | Cat# ab80522, RRID:AB_1603675 |
| Rabbit monoclonal anti-BMP2 | Abcam | Cat# ab214821; RRID:N/A |
| Mouse monoclonal anti-PCNA | Santa Cruz | Cat# sc-25280, RRID:AB_628109 |
| Mouse monoclonal anti-SOX18 | Santa Cruz | Cat# sc-166025, RRID:AB_2195662 |
| Rat monoclonal anti-PRDM1 | Thermo Fisher | Cat# 14-5963-80, RRID:AB_1907438 |
| Rabbit polyclonal anti-SOX9 | Millipore | Cat# AB5535, RRID:AB_2239761 |
| Donkey Anti-Rabbit IgG H&L (Alexa Fluor® 555) | Abcam | Cat# ab24273; RRID:AB_778767 |
| Goat Anti-Mouse IgG H&L (Alexa Fluor® 488) | Abcam | Cat# ab150113; RRID:AB_2576208 |
| HRP-labeled Goat Anti-Rat IgG(H+L) | Beyotime | Cat# A0192; RRID:NA |
| HRP-labeled Goat Anti-Rabbit IgG(H+L) | Beyotime | Cat# A0208; RRID:NA |
| HRP-labeled Goat Anti-Mouse IgG(H+L) | Beyotime | Cat# A0216; RRID:NA |

The next morning, the slides were further incubated with secondary antibodies at 37 °C for 30 min. DAPI was used to stain nuclei and the slides were mounted with anti-fade mounting medium. Pictures were taken under LEICA TCS SP5 II confocal microscopy (Leica Microsystems GmbH, Wetzlar, Germany). For enzyme substrate-based immunohistochemistry, the slides were washed with 3 % H_2_O_2_ for 10 min to block endogenous peroxidase activity prior to blocking and DAB (ZSGB-BIO, Bejing, China) solution was used for chromogenic reaction.

### Single cell suspension preparation

For each group skin tissues were obtained from at least 6 independent male fetuses prior to digestion. To prepare dorsal skin single cell suspension for single-cell RNA sequencing, the fetus back skin tissues were isolated via microdissection and 0.25 % trypsin/EDTA solution was used to digest E13.5 and E16.5 fetus dorsal skin tissues at 37 °C for 5 min. For 0 dpp mice dorsal skin tissues, 2 mg/ml collagenase IV (Sigma, St Louis, MO, USA) was used to digest skin tissues at 37 °C for 30 min. After trypsinization, the skin tissues were mechanically dissociated into single cell suspension by pipetting, the cell suspensions were then filtered through a 40 μm nylon cell strainer (BD Falcon, BD Biosciences, San Jose, CA, USA) prior to single cell library construction.

### Single cell library preparation and sequencing

Single cell barcoding and library preparation were performed based on 10x Genomics single-cell RNA sequencing platform (10x Genomics, Pleasanton, CA, USA). Briefly, the single cell suspension prepared above were immediately counted using a hemocytometer (TC20, Bio-Rad, Hercules, CA, USA) and the cell concentrations were adjusted to 1000 cells/μl prior to barcoding. To barcode the single cells with 10x Barcoded gel beads, 10x Genomics Chromium Single Cell 3’ Library & Gel Bead Kit v2 (10x Genomics Inc., Pleasanton, CA, USA, 120237) and 10x Genomics Chromium barcoding system was used to construct 10x barcoded cDNA library following the manufacturer’s instructions. Illumina HiSeq X Ten sequencer (Illumina, San Diego, CA, USA) was used for sequencing and pair-ended 150 bp (PE150) reads were generated for downstream analysis.

### 10x sequencing data preprocessing

The CellRanger (v2.2.0) software was used for analyzing raw sequencing data according to 10x Genomics official pipeline (https://support.10xgenomics.com/single-cell-gene-expression/software/pipelines/latest/what-is-cell-ranger). Briefly, the sequencing raw base call (BCL) files were firstly transformed into FASTQ files with ‘cellranger mkfastq’ function. The generated FASTQ files were then processed with ‘cell ranger count’ wrapped function with ‘--force-cells = 7000’ argument to adjust sample size. Cell ranger count function used wrapped STAR software to align sequence to the reference genome and the 10x pre-built mouse genome (mm10-3.0.0) was used (https://support.10xgenomics.com/single-cell-gene-expression/software/downloads/latest). The output files containing gene expression matrices and barcode information of CellRanger pipeline were then used for downstream visualization analysis.

### Characterization of cell clusters

After CellRanger pipeline, the quality control (QC) and cell clustering were analyzed with single-cell RNA seq Seurat software (v2.3.4) based on R environment (R version: 3.5.1, https://www.r-project.org/) following the online guide (https://satijalab.org/seurat/). We used ‘filtered_gene_bc_matrices’ files generated by CellRanger as input files for Seurat. For each dataset, we firstly filtered cells with unique detected genes less than 200 and genes detected less than 3 cells, then we used ‘FilterCells’ function to remove cells with a total number of detected genes (nGenes) less than 1750. After normalization, the variable genes for each dataset were calculated for downstream clustering assay.

To compare transcriptome profiles along three different developmental timepoints, we then merged three different datasets using ‘RunMultiCCA’ function implemented in Seurat. RunMultiCCA used a canonical correlation analysis to remove variation caused by sample source. After dataset alignment, we then performed a clustering analysis on the integrated dataset based on t-distributed Stochastic Neighbor Embedding (tSNE) algorithm implemented in Seurat. To identify cluster specifically expressed genes, we used Seurat implemented ‘FindAllMarkers’ function to calculate cluster markers and the tSNE identified cell clusters were annotated with based on the previously reported canonical marker genes expression.

To subcluster cell clusters of interest for in-depth analysis and/or downstream differentiation trajectory construction, we used Seurat implemented ‘SubsetData’ function to extract cluster of interest. The extracted subclusters were then re-run the Seurat pipeline, which provides higher resolution for dissecting cellular heterogeneity among particular cell types.

### Constructing single cell pseudotime differentiation trajectory

To interpret cell differentiation fate decisions, we used Monocle (v 2.10.0) to order single cells along pseudotime according to the official tutorial (http://cole-trapnell-lab.github.io/monocle-release/docs/#constructing-single-cell-trajectories). To perform pseudotime ordering to particular cell types, we firstly subclustered interested cell type from Seurat object, then, the Monocle object was constructed using ‘newCellDataSet’ function in Monocle. To order single cells along pseudotime, we used Seurat identified variable genes as ordering genes to construct single cell differentiation trajectory. The root state was set according to cell Seurat identified cell cluster label and ‘BEAM’ function was used to calculate branch-specific expressed genes. To plot branch-specific expression heatmap, we used Monocle implemented ‘plot_genes_branched_heatmap’ function and genes with qval < 1e-4 were regarded as input genes. Gene clusters were further divided into four clusters according to k-means. To investigate gene functions in each gene clusters, we used Metascape (http://metascape.org/gp/index.html#/main/step1) to perform gene ontology (GO) analysis.

### Core TFs prediction

To infer core TFs within each branch identified by Monocle, we used SCENIC algorithm to infer regulon activity within each cell states. The analysis pipeline was performed following the developer’s instructions (https://github.com/aertslab/SCENIC). Briefly, the expression matrix was firstly extracted from Seurat and was further transformed into SCENIC required format (rows represent genes, columns represent cells) in R, we then extracted cellular branch information identified by Monocle to construct expression matrix of desired states.

We further filtered cells with genes that were detected in less than 1% of the total cells and genes with less than at least 6 UMI counts across all samples. Then, GENIE3 was used to identify co-expressed gene modules and infer potential TF targets for each module based on the expression matrix. After that, RcisTarget was used to perform cis-regulatory motif analysis, we scanned two motif to TFs databases (mm10 refseq-r80 10kb_up_and_down_tss and mm10 refseq-r80 500bp_up_and_100bp_down_tss; https://resources.aertslab.org/cistarget/) and kept modules with significant motif enrichment, this modules were then termed as regulons according to SCENIC pipeline. To visualize regulon activity within each cellular state identified by Monocle, we further used SCENIC pipeline to binarize the regulon network activity based on AUCell algorithm, and we used binary regulon activity matrix to visualize regulon activity within each cellular state.

### Cell to cell ligand-receptor interaction analysis

To infer the hypothetical intercellular communication, we compared cell type-specific DEGs identified by Monocle and manually sorted ligand-receptor pairs according to cell relationship. The mouse ligand-receptor pairs used here were compiled by Daniel et al and about 2,009 ligand-receptor pairs were used in this study. After sorting ligand-receptor pairs, we used igraph (https://github.com/igraph/rigraph) and edgebundleR (https://github.com/garthtarr/edgebundleR) R packages to visualize the ligand-receptor networks and the node represents genes while the solid line connects each ligand-receptor pair. To avoid confusion, we separately plotted E13.5 to E16.5 and E16.5 to P0 ligand-receptor pairs. Only cell types at the same time point and shares differentiation relationship were considered to sort ligand-receptor pairs.

### Data availability

The single cell RNA sequencing data used in this research is deposited in NCBI GEO databases under accession number: GSE131498.

## Acknowledgments

This work was supported by National Nature Science Foundation (31671554 & 31772573) and National Key Research and Development Program of China (2018YFC1003400). The authors would like to thank Dr. Paul Dyce at Auburn University for his careful editing of this manuscript.

## Supplementary Figures

**Figure S1: Quality control of single-cell data.** (A) Single cell datasets quality metrics summary identified by CellRanger. (B) Violin plot displaying the number of genes (nGene), UMI (nUMI), and percentage of mitochondrial genes (percent.mito) detected in all single cells from three different datasets. The gene to UMI relationship for each dataset was also visualized. Generally, the more UMI captured, the more genes detected.

**Figure S2: Characterization of major cell populations in the embryonic skin.** (A) Hierarchical clustering of different cell clusters identified by tSNE. the cluster number was in accordance with Figure 1B. (B) Visualization of canonical maker gene expression in the tSNE plot of all single cells. DC markers: *Lef1*, *Prdm1*, *Sox18*, *Inhba*; Endothelial markers: *Pecam1*, *Kdr*; Melanocyte markers: *Plp1*, *Fabp7*; Pericyte markers: *Rgs5*, *Acta2*: Neural markers: *Map2*, *Stmn3*: Muscle markers: *Myod*, *Pax7*, Immune markers: *Cd52*, *Fcer1g*. (C) Heatmap displaying top 10 signature gene expression in each cluster.

**Figure S3: Visualization of key DC markers expression.** (A) Single cell pseudotime trajectory color-coded by pseudotime. (B) Visualization of representative gene set specific gene expression projected in the pseudotime trajectory.

**Figure S4: Investigating gene expression profile during AA/ZZ DP and G-DP fate commitment** (A) Gene expression profile and representative genes during AA/ZZ DP and G-DP fate commitment. (B) Representative AA/ZZ DP and G-DP marker expression projected into pseudotime trajectory. (C) Comparison of Fgf7, Lef1, Gli1, and Notch1 expression in the pseudotime trajectory. The red dotted box depicts an intermediate stage, while the blue line indicates the endpoint. (D) Circos plot indicating the shared GO terms and heatmap of top GO comparison among 4 different gene sets. Genes belong to the same GO were connected together in the circus plot.

**Figure S5: Interpreting HFSC and matrix molecular signature** (A) Scatter plot of mean expression against the empirical dispersion displaying the variable genes (black dots) used for Monocle pseudotime ordering. (B) Pseudotime ordering analysis of all epithelium populations, cells were labeled with pseudotime (left) and their corresponding cluster information (right), respectively. (C) HFSC and matrix precursor marker genes expression along pseudotime. Cells were labeled with cell state corresponding to figure 4B. (D) Visualization of *Dkk4* and *Lef1* expression projected into pseudotime trajectory.

**Figure S6: Dissecting hair shaft and IRS fate decisions.** (A) Hair shaft and IRS signature gene expression along pseudotime. Cells were labeled with cell states with a solid line indicating fate 2 and a dotted line indicating fate 1. (B) Comparison of HS and IRS signature genes between Anagen VI from Yang et al., and our single cell analysis here. The overlapped genes were listed in the rectangular box.

## Supplementary Tables

**Supplementary Table 1.** List of DEGs for tSNE identified 13 clusters. Corresponding to Figure 1B.

**Supplementary Table 2.** List of branch-specific DEGs expression along pseudotime. Corresponding to Figure 2C.

**Supplementary Table 3.** List of SCENIC enriched TFs and it’s corresponding targets.

Supplementary Table 4. Signature genes comparison P0 G-DP, AA/ZZ DP (this study) and AA/ZZ DP, ZZ DP, GAA DP, G-DP (Rezza et al.). Corresponding to Figure 3D.

**Supplementary Table 5.** Signature genes comparison E16.5 matrix/HFSC precursors (this study) and P5 matrix/HFSC (Rezza et al. 2016). Corresponding to Figure 4D.

**Supplementary Table 6.** Signature genes comparison P0 hair shaft/IRS (this study) and anagen VI HS/IRS (Yang et al. 2017).

