## Supplementary Figures 1-6 for "Single-cell transcriptome profiling reveals dermal and epithelium cell fate decisions during embryonic hair follicle development"

### Supplementary Figure 1

(A)

| Sample info. | E13.5 | E16.5 | E19.5 |
|---|---|---|---|
| Estimated Number of Cells | 7,000 | 7,000 | 7,000 |
| Valid Barcodes | 97.2% | 97.3% | 97.6% |
| Mean Reads per Cell | 75,538 | 77,370 | 75,560 |
| Median Genes per Cell | 2,434 | 2,978 | 1,989 |
| Total Genes Detected | 19,997 | 19,767 | 19,145 |
| Reads Mapped to Genome | 90.5% | 91.5% | 94.0% |
| Reads Mapped Confidently to Transcriptome | 62.8% | 65.2% | 71.7% |

(B)

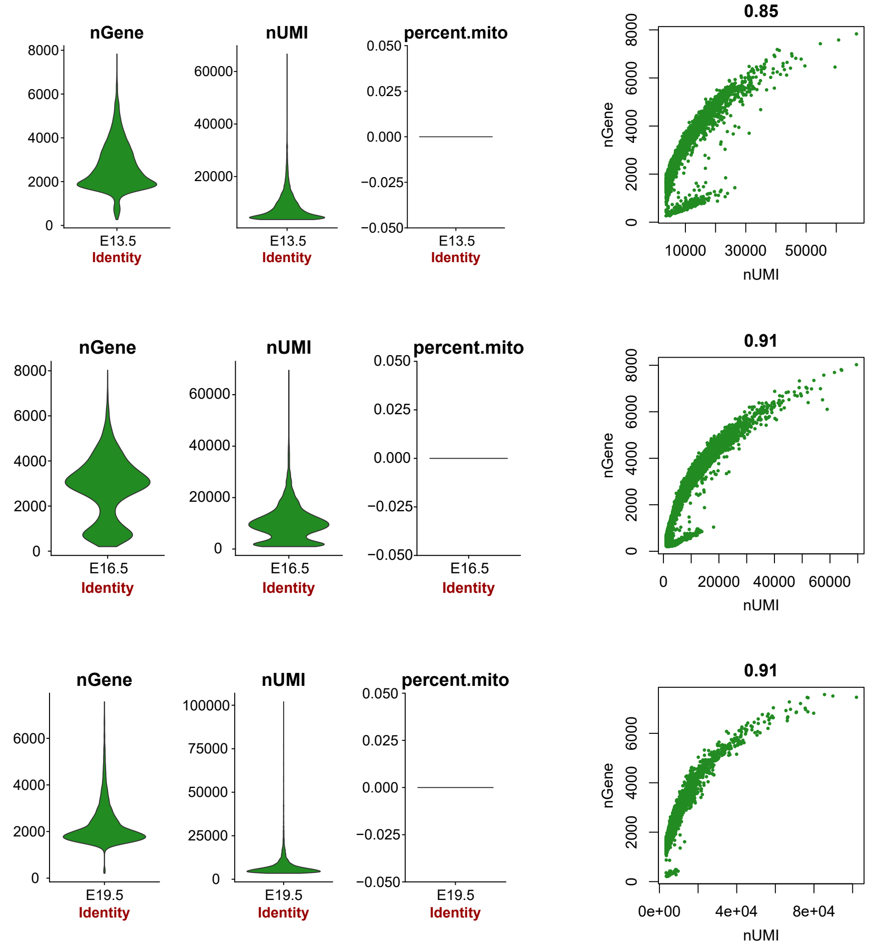

### Supplementary Figure 2

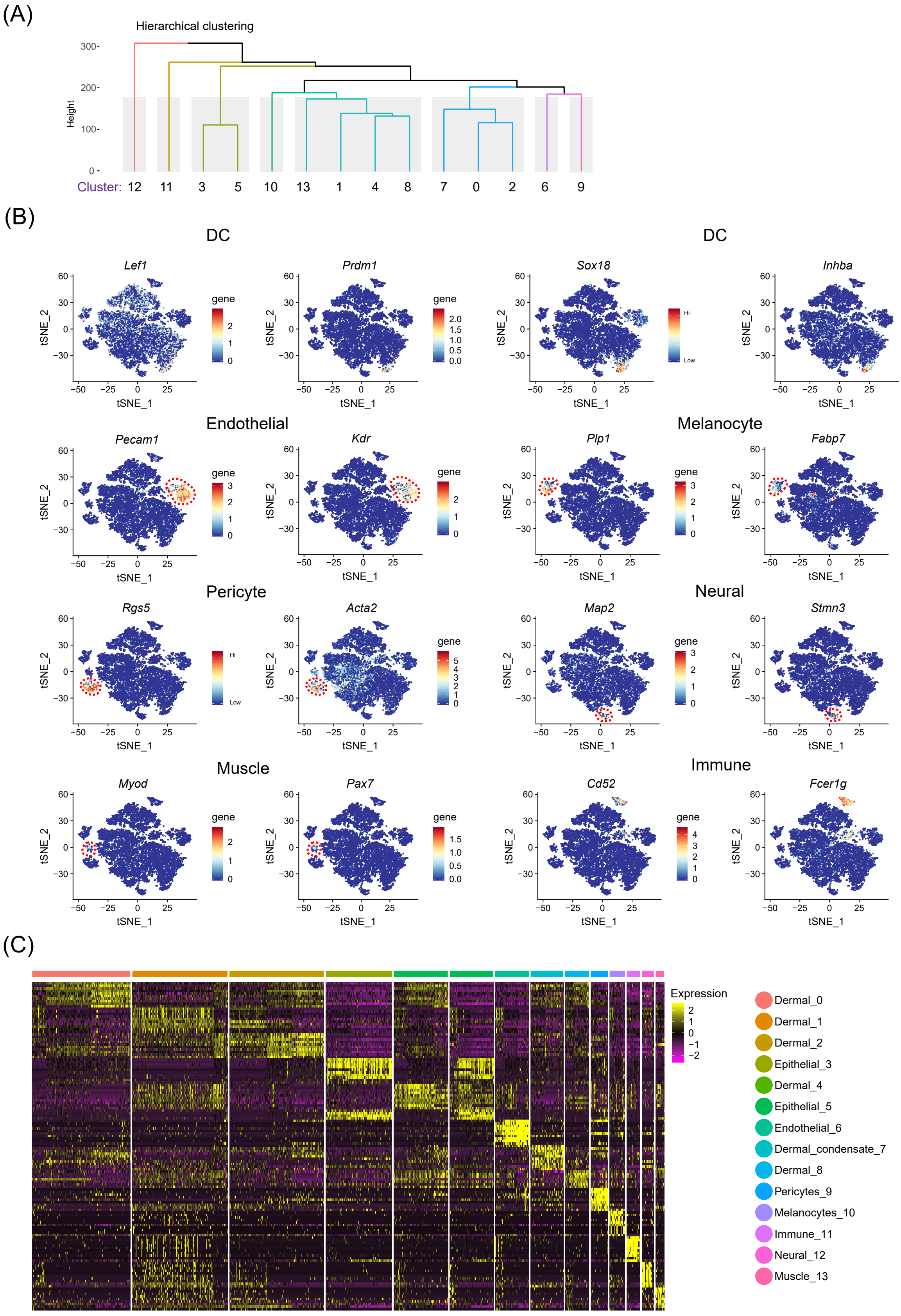

### Supplementary Figure 3

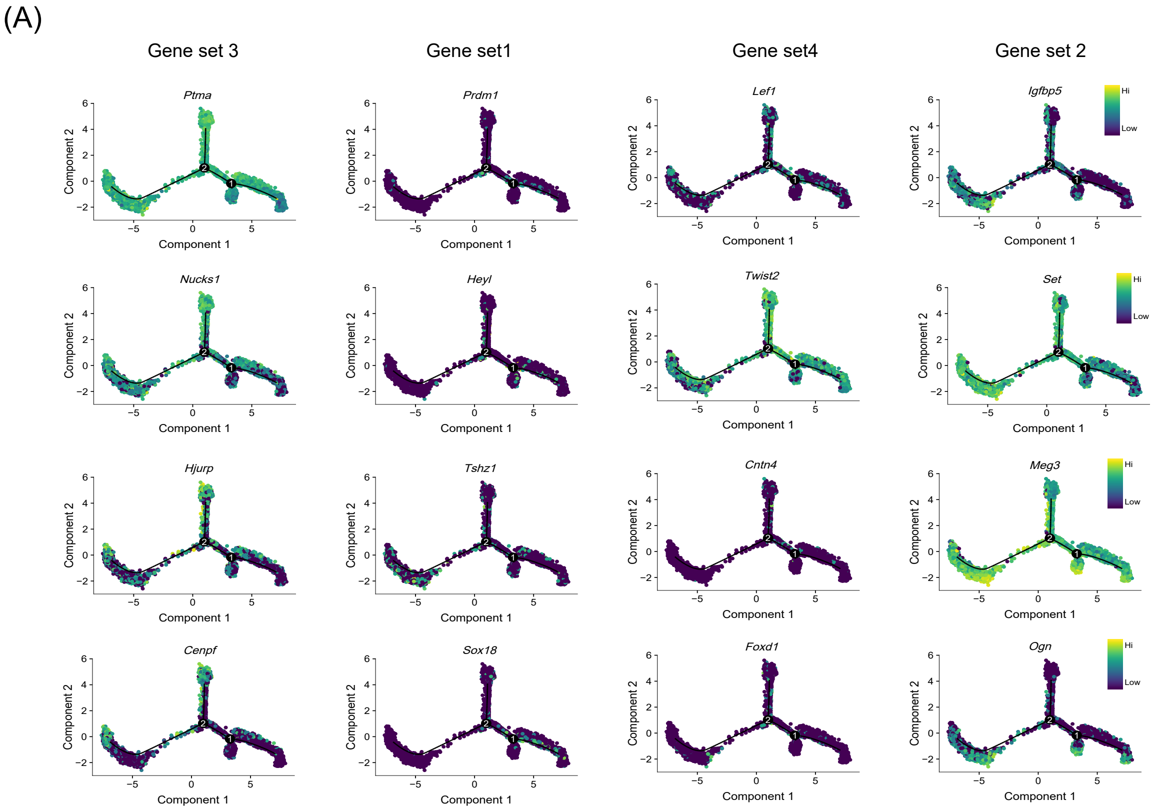


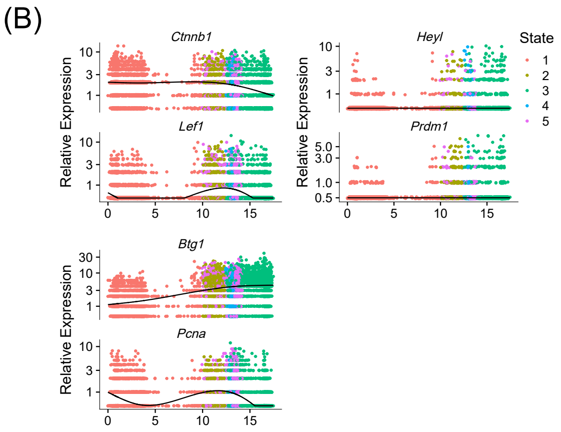


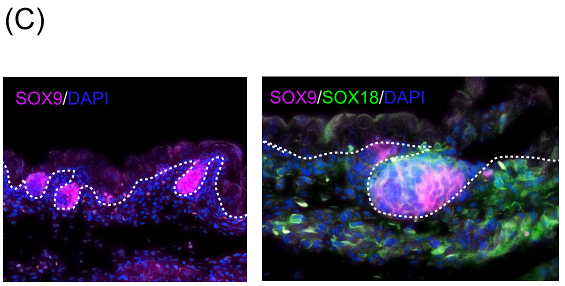

### Supplementary Figure 4

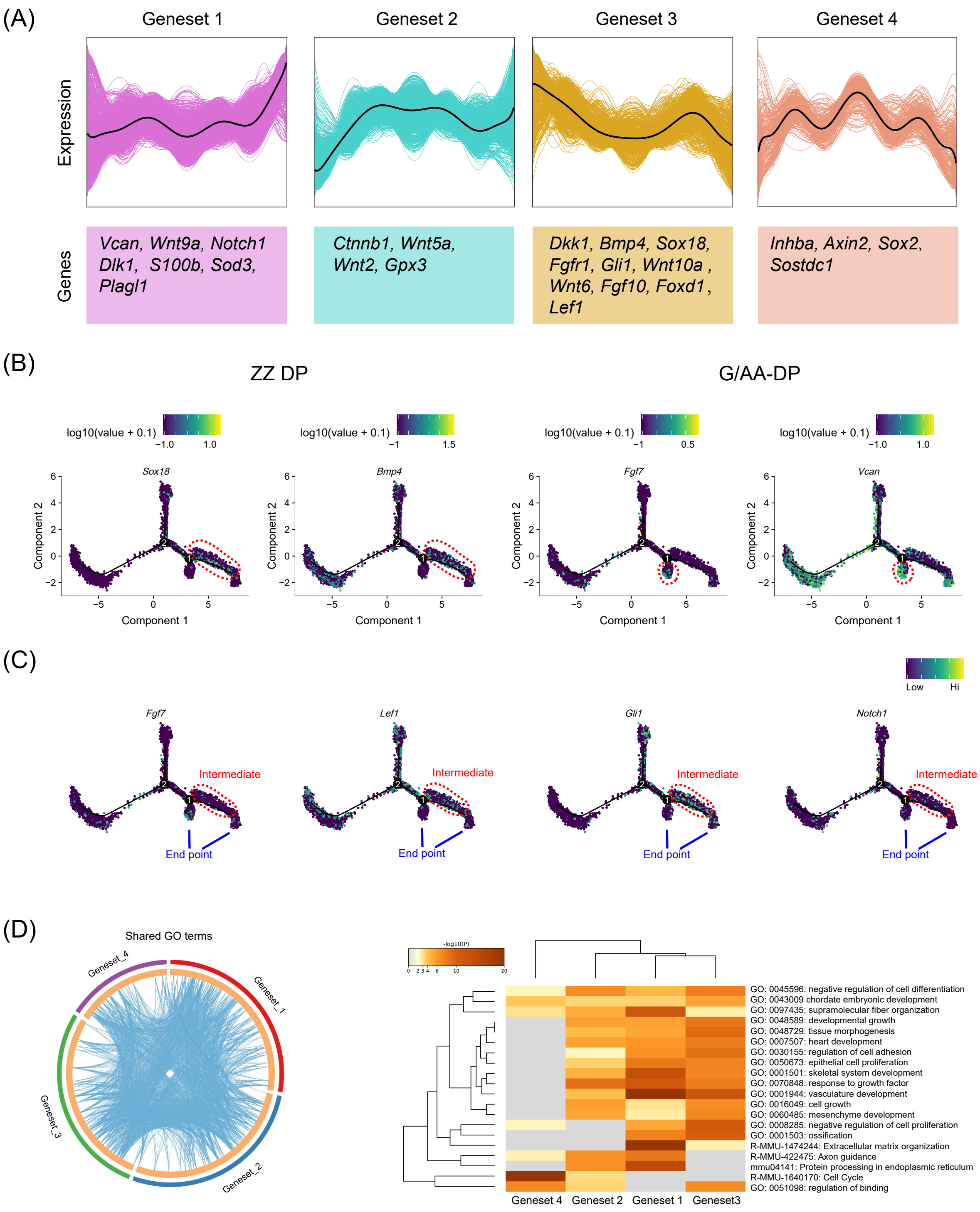

### Supplementary Figure 5

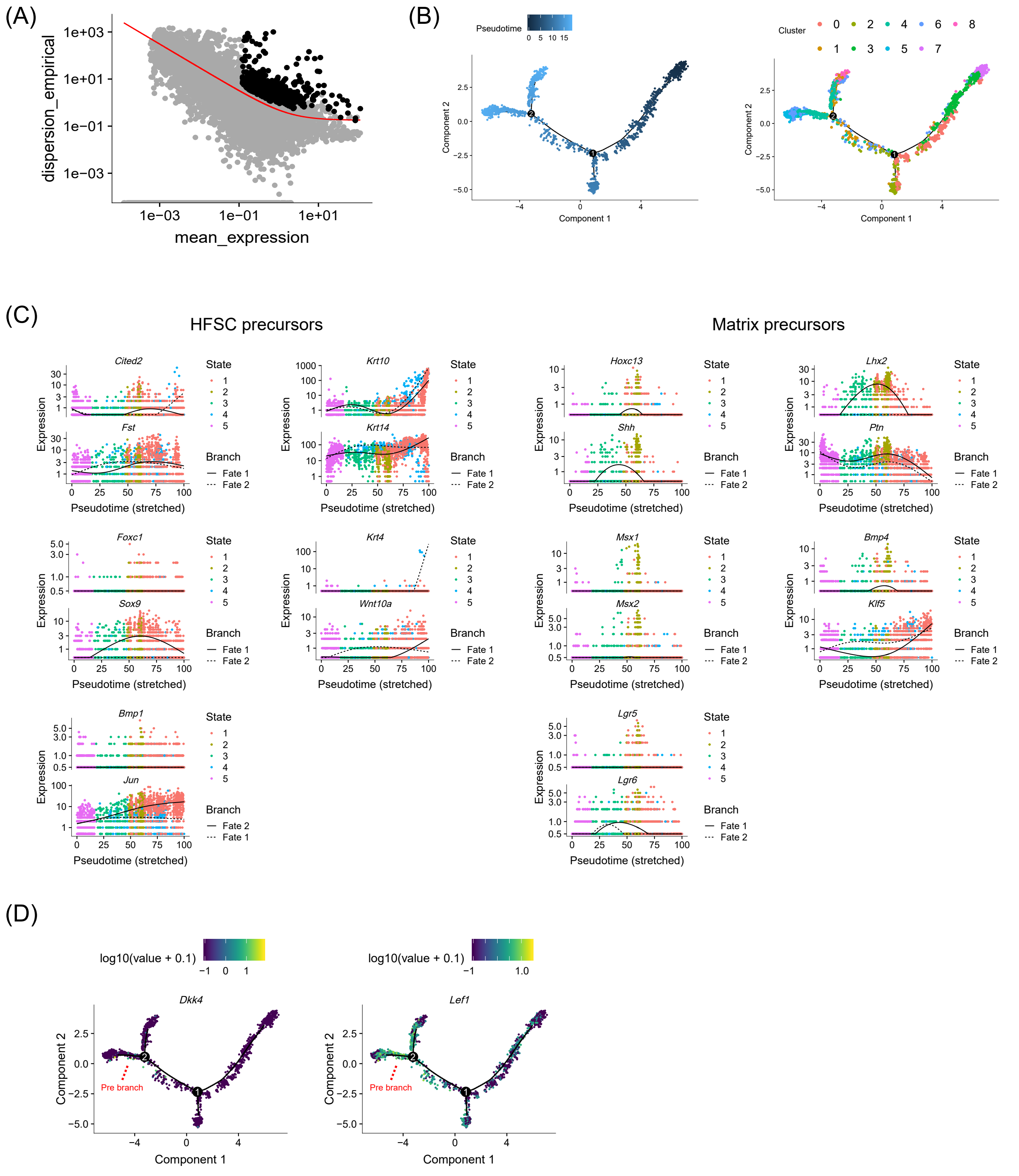

### Supplementary Figure 6

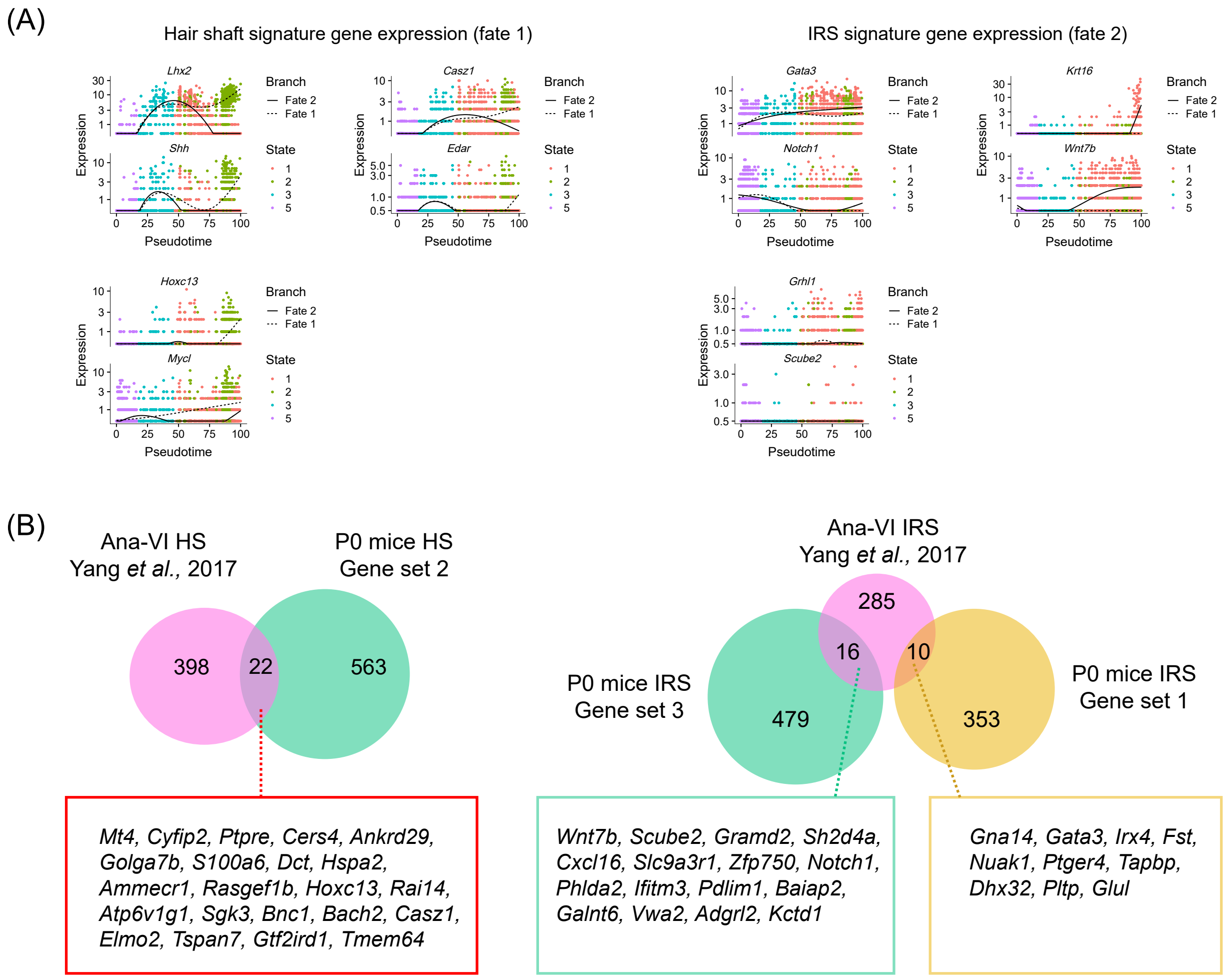
